# Effective QTL Discovery Incorporating Genomic Annotations

**DOI:** 10.1101/032003

**Authors:** Xiaoquan Wen

## Abstract

Mapping molecular QTLs has emerged as an important tool for understanding the genetic basis of cell functions. With the increasing availability of functional genomic data, it is natural to incorporate genomic annotations into QTL discovery. In this paper, we describe a novel method, named TORUS, for integrative QTL discovery. Using hierarchical modeling, our approach embeds a rigorous enrichment analysis to quantify the enrichment level of each annotation in target QTLs. This enrichment information is then used to identify QTLs by up-weighting the genetic variants with relevant annotations using a Bayesian false discovery rate control procedure. Our proposed method only requires summary-level statistics and is highly efficient computationally: it runs one-hundred times faster than the current gold-standard QTL discovery approach that relies on permutations. Through simulation studies, we demonstrate that the proposed method performs accurate enrichment analysis and controls the desired type I error rate while greatly improving the power of QTL discovery when incorporating informative annotations. Finally, we analyze the recently released expression-genotype data from 44 human tissues generated by the GTEx project. By integrating the simple annotation of SNP distance to transcription start sites, we discover more genes that harbor expression-associated SNPs in all 44 tissues, with an average increase of 1,485 genes.

## 1 Background

With the advancements in sequencing technology, mapping quantitative trait loci (QTL) with cellular phenotypes has emerged as a powerful tool for understanding the genetic basis of cell functions. Recent QTL mapping studies using RNA-seq, ChIP-seq, DNaseI-seq, ATAC-seq and DNA methylation data have revealed that an abundance of genetic variants are associated with various cellular phenotypes (Lappalainen *et al*. 2013; Ardlie *et al*. 2015; Ding *et al*. 2014; Degner *et al*. 2012; McVicker *et al*. 2013; Banovich *et al*. 2014). Subsequently, the discovery of molecular QTLs has provided valuable insights for understanding the molecular mechanisms of complex diseases (Albert and Kruglyak 2015). Note that a distinctive feature of molecular QTL analysis is that tens of thousands of molecular phenotypes are simultaneously measured (e.g., genome-wide gene expression profiling by RNA-seq), which imposes a new type of statistical challenge.

In this paper, we use the term QTL to refer to the genomic regions that harbor traits associated causal variants, and following Veyrieras *et al*. (2008), we refer to the actual causal variants as quantitative trait nucleotides (QTNs). In practice, the statistical analysis of molecular QTLs typically consists of two stages: the primary goal of the first stage is to screen a large number of candidate loci and identify QTLs, and we refer to this process as *QTL discovery;* in the second stage, a fine-mapping analysis is performed to determine the potential QTNs in each discovered QTL. In addition to providing a list of candidate QTLs for fine-mapping analysis, QTL discovery is also highly important for network and pathway analysis. When analyzing expression quantitative trait loci (eQTLs), the candidate genomic regions are generally formed by the coding and the neighboring regulatory regions of each target gene, and the QTL discovery analysis is also known as eGene discovery (Lappalainen *et al*. 2013; Ardlie *et al*. 2015; Sul *et al*. 2015). The statistical strategies for the two stages of QTL analysis differ: the QTL discovery is typically formulated as a multiple hypothesis testing problem, whereas the QTN fine mapping is often addressed using variable selection techniques.

Our primary focus in this paper is the statistical analysis for QTL discovery. The gold-standard QTL discovery approach has been well established in *cis*-eQTL analysis and can be trivially generalized to other molecular QTL analyses (Flutre *et al*. 2013; Lappalainen *et al*. 2013; Ardlie *et al*. 2015). In *cis*-eQTL mapping, the standard approach first performs single SNP association testing for all gene-SNP pairs. For each gene, the minimum *p*-value from all member SNPs is then regarded as the gene-level test statistic and is subsequently converted into a gene-level *p*-value to test the null hypothesis asserting no associated SNPs within the locus of interest. Because the null distribution of the gene-level test statistic is generally unknown due to complicated linkage disequilibrium (LD) patterns, extensive permutations are required to obtain the gene-level *p*-values. Finally, false discovery rate (FDR) control procedures, e.g., Benjamini-Hochberg (Benjamini and Hochberg 1995) and Storey’s *q*-value procedure (Storey 2003), are applied to correct for multiple testing of tens of thousands of genes genome-wide.

Although the gold-standard procedure is statistically justified and widely applied, it lacks computational efficiency due to its heavy reliance on extensive permutations: the computational cost remains very high for genome-wide QTL discovery, even though recent work (Sul *et al*. 2015) has made significant improvements. More critically, there is no principled way to flexibly incorporate valuable genomic annotations into the standard procedure. With the increasing availability of functional genomic data (Pique-Regi *et al*. 2011; ENCODE Project Consortium *et al*. 2012; Kundaje *et al*. 2015), the scientific community has accumulated substantial knowledge on the functional roles of individual genetic variants. It is completely intuitive to incorporate this knowledge into the analysis of QTL discovery and prioritize the genomic loci that harbor well-annotated functional variants. Similarly, existing results regarding QTL analyses can also provide valuable insights into the distributive patterns of causal QTNs. For example, almost all of the analyses in *cis*-eQTL mapping report that associated casual SNPs tend to cluster around transcription start sites (TSS) and that the abundance of signals rapidly decreases away from TSS (Lappalainen *et al*. 2013; Wen *et al*. 2015; Ardlie *et al*. 2015). In light of this pattern, it appears natural to up-weight the SNPs close to TSS in eQTL analysis *a priori* rather than treating every *cis*-SNP equally. However, to the best of our knowledge, a principled approach that can effectively incorporate known genomic annotations into the analysis of QTL discovery does not exist.

In this paper, we use a natural hierarchical model to integrate SNP-level annotations into QTL mapping. We propose a highly efficient computational framework, TORUS, to first evaluate the enrichment level of each genomic annotation in the causal variants, and then we utilize the quantified enrichment information to prioritize each candidate SNP accordingly and perform Bayesian multiple hypothesis testing for QTL discovery. Importantly, the proposed QTL discovery approach also seamlessly connects to our proposed Bayesian QTL fine-mapping method (Wen 2014; Wen *et al*. 2015) by providing the SNP-level prior information as a by-product. Through simulations, we demonstrate the superiority of the proposed approach over the state-of-the-art gold-standard approach in terms of both computational efficiency and power of QTL discovery. Finally, we demonstrate our approach by analyzing eQTL data from 44 human tissues that were recently released by the GTEx project.

## 2 Results

### 2.1 Method Overview

Our statistical approach is built upon our recently proposed Bayesian hierarchical model (Wen *et al*. 2015). A distinctive feature of this model is its use of prior specification to quantitatively connect the genomic annotations of a candidate SNP with its latent trait-association status through a set of regression coefficients known as *enrichment parameters* (details are shown in the Methods section). This model has been successfully applied in fine-mapping eQTNs across multiple tissues (Ardlie *et al*. 2015) and across multiple populations (Wen *et al*. 2015). Moreover, this model naturally accounts for LD between candidate SNPs and shows great advantages over the existing standard single-SNP-based QTL analysis approaches. In this paper, we extend this approach for QTL discovery applications.

For each genomic locus *l* consisting of *p* SNPs, we use a binary *p*-vector *γ_l_* to denote the latent association status of all member SNPs, i.e., 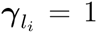 indicates that the *i*-th SNP is associated with the quantitative trait of interest. The problem of QTL discovery can then be formulated as testing the null hypothesis *H*_0_: *γ_l_* = 0 for each locus. When mapping molecular QTLs, tens of thousands of loci are simultaneously interrogated; therefore, the issue of multiple testing control/correction must be addressed.

To take full advantage of the Bayesian hierarchical model, we employ a Bayesian false discovery rate (FDR) control approach (Newton *et al*. 2004; Müller *et al*. 2004). Briefly, this procedure requires computing the posterior probability

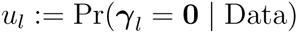

to summarize the evidence for (or against) the null hypothesis for each locus *l*. The null hypothesis is intuitively rejected if the corresponding *u_l_* is too small. To determine the rejection threshold *t_α_* based on a predefined FDR control level *α* (typically set at 0.05), a straightforward algorithm (Newton *et al*. 2004) can be applied such that the expected value of false discovery proportion (which can be directly computed from *u_l_*s) from the observed data is not greater than *α*. The connection between the Bayesian FDR control approach and commonly applied frequentist approaches has been well documented in the statistical literature (Newton *et al*. 2004; Müller *et al*. 2004), and we also provide a brief account in the Methods section and in the Supplementary Material.

By computing *u_l_* based on the proposed Bayesian hierarchical model, the Bayesian FDR control procedure naturally allows genomic annotations to be leveraged in QTL discovery. However, the exact evaluation of *u_l_* requires integrating out all enrichment parameters and exploring an enormous space of all possible association models. This becomes a computationally daunting challenge even for a single locus, let alone the genome-wide application for tens of thousands of loci. To overcome the computational difficulty, we propose evaluating *u_l_* using an approximation *û_l_* that can be computed considerably more efficiently. Specifically, rather than fully integrating out the enrichment parameter ***α***, we apply an EM algorithm to determine its maximum likelihood estimate (MLE), 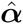, and evaluate an empirical Bayes estimate of the posterior probability, namely, 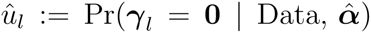. In addition, to reduce the extensive exploration of all possible values of *γ_l_*, we adopt a strategy that focuses only on a subset of alternative models that contain at most K QTNs and that assumes that the posterior probabilities of association models with more than K QTNs are negligible. Because the convincing QTNs discovered from the association data are highly *sparse* compared with the number of candidate SNPs in almost all genetic/genomic applications, this approximation is likely accurate for the vast majority of loci. For the small number of loci that do harbor more than K casual QTNs, this specific approximation leads to a conservative over-estimation of *u_l_*, which may result in a loss of power but no inflation of type I error. To achieve the best computational efficiency, in practice, we set *K* = 1, which essentially assumes at most one QTN per locus. It can be argued that the gold-standard frequentist approach also implicitly makes such an assumption (see the Methods section). Furthermore, although this very assumption has been successfully applied by other Bayesian approaches (Pickrell 2014; Servin and Stephens 2007; Veyrieras *et al*. 2008; Flutre *et al*. 2013), it has always been formulated as an explicit *prior* assumption and hence requires a somewhat non-natural parameterization that is not only difficult to interpret but also complicates the estimation of the enrichment parameters. Jointly applying both approximation strategies results in a highly efficient computational procedure for QTL discovery. Moreover, the proposed approach only requires summary-level statistics from single SNP association testing, which is extremely convenient for addressing genome-wide high-throughput genomic data.

The statistical methods are implemented in the software package TORUS, which is freely available at https://gihub.com/xqwen/torus/.

### 2.2 Simulation Study

We perform a series of simulation studies to demonstrate the power, robustness and computational efficiency of the proposed QTL discovery approach.

#### 2.2.1 QTL Discovery without Annotation

In the first simulation, we generate genome-wide eQTL data sets assuming no influence from any genomic feature. Our goal is to evaluate the performance of the proposed Bayesian procedure under the baseline scenario and to compare it with the commonly applied gold-standard approach.

We select 11,761 protein coding and linc-RNA genes from the GEUVADIS project (Lappalainen *et al*. 2013) and the genotype data from 343 European individuals. For each gene, we randomly select 50 *cis*-SNPs with a minor allele frequency of ≥ 0.05. With probability 1 − *π*_0_, we randomly assign 1 to 3 eQTNs. Given the eQTNs for each gene, we simulate the expression levels using a multiple regression model (Supplementary Materials). We generate 20 data sets for each *π*_0_ value and vary the value of *π*_0_ from 0.1 to 0.9.

Without using annotation information, the Bayesian hierarchical model is reduced to a simple form with a single parameter in the logistic prior, which assumes *a priori* each candidate SNP independently and equally likely to be the causal eQTL. For comparison, we analyze the simulated data sets using the software package eGENE-MVN (Sul *et al*. 2015). This package implements the gold-standard QTL discovery method that uses the minimum single SNP association *p*-value in a locus as the test statistic; however, it finds the corresponding locus-level *p*-value in a considerably more efficient manner. After obtaining the locus-level *p*-values, we perform FDR control and identify QTLs using Storey’s *q*-value method.

The simulation results (Table 1) indicate that both TORUS and the gold-standard approach control FDR at the desired level across all *π*_0_ levels. Overall, the powers of the two methods are quite similar. However, the standard method achieves slightly better power when *π*_0_ is small, and the proposed Bayesian method exhibits better performance when *π*_0_ is relatively large. We suspect that this result is mainly because when *π*_0_ is large, the “one causal SNP per locus” assumption becomes closer to the truth compared with the situation when *π*_0_ is small, and therefore, the posterior probability approximation is more accurate and less conservative. In addition, to examine the robustness of the proposed approach, we re-analyze the simulated data but include the annotation of SNP distance to TSS (details described in simulation study II).

**Table 1:** Comparison of TORUS and the gold-standard minimum *p*-value-based QTL discovery procedure without genomic annotations using simulated eQTL data. Both methods control the desired FDR level at 5%. The standard method performs better when the proportion of candidate loci being QTLs (i.e., 1 − *π*_0_) is high, whereas the proposed procedure works better when the proportion is low. In all cases, however, the powers are comparable. In addition, we perform the proposed approach using SNP distance to TSS as an annotation, and the results remain virtually identical, as expected.

| $\pi_0$ | TORUS without annotation | | Minimum $p$-value Method | |
| --- | --- | --- | --- | --- |
|  | FDR | Power | FDR | Power |
| 0.10 | 0.010 | 0.842 | 0.024 | 0.864 |
| 0.33 | 0.028 | 0.807 | 0.038 | 0.810 |
| 0.50 | 0.038 | 0.801 | 0.040 | 0.789 |
| 0.67 | 0.045 | 0.767 | 0.047 | 0.743 |
| 0.90 | 0.049 | 0.739 | 0.049 | 0.701 |

As expected, the enrichment analysis indicates that there is little impact of the annotation to the eQTLs in the simulated data set (due to our simulation scheme), and the results for eGene discovery remain virtually identical.

Most importantly, our computational time benchmark shows that the proposed Bayesian method is considerably more efficient: to analyze a single simulated data set on a Linux box with an 8-core Intel Xeon 2.13 GHz CPU, the average running time for the Bayesian method is approximately 2 minutes 25 seconds (with 12 parallel processing threads); in comparison, eGENE-MVN requires approximately 3 hours and 45 minutes (also with 12 parallel threads) for the same computational task.

#### 2.2.2 QTL Discovery with Annotation

Our second simulation study attempts to mimic a commonly observed phenomenon in *cis*-eQTL mapping: eQTL signals tend to cluster around transcription start sites of the corresponding target genes and rapidly decrease away from TSS. We use the same set of 11,761 genes from the GUEVADIS project but include all SNPs within a 1 Mb radius from the TSS of each gene. On average, there are 5,856 SNPs per gene (median of 5,892). We do not impose any restrictions on the minor allele frequencies of the *cis*-SNPs and take all the genotypes directly from the GUEVADIS project. During the simulation, causal eQTL SNPs are randomly assigned by a probability computed from a continuous function of SNP distance to TSS (DTSS, measured in kb and denoted by *d*), i.e.,

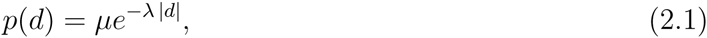

where *λ* controls the rate of decay in the expected number of causal eQTL SNPs away from TSS and *μ* determines the overall expected number of *cis*-eQTLs. We experiment with two different *λ* values, *λ* = 0.02 and *λ* = 0.1, corresponding to relatively modest and fast rates of decay, respectively. We then set the *μ* values to keep the overall expected number of causal eQTL SNPs comparable across the schemes. Note that our simulation function (2.1) is not compatible with the functional form of our logistic prior (4.2).

For each simulation setting, we generate 20 data sets and analyze each data set with and without incorporating DTSS information. We do not run the minimum *p*-value-based approach on these considerably larger data sets because of the high computational cost. Nevertheless, we fully expect its performance to be similar to that of the proposed Bayesian approach ignoring DTSS information based on our evaluation in the first simulation study. When utilizing DTSS information, we follow the approaches used in Veyrieras *et al*. (2008); Degner *et al*. (2012) to dissect the genome into variable sizes of distance bins. In general, we use smaller sized bins in the close vicinity of TSS and larger sized bins away from TSS. The details on the binning of SNPs are given in the Supplementary Materials.

Our results indicate that by estimating the enrichment parameters, the Bayesian approach effectively characterizes the impact of the DTSS on the eQTL abundance. The estimation of the eQTL signal decay rates with respect to TSS is quite accurate (Fig. 1), albeit our estimation model is very different than the data generative model. We also find that the baseline prevalence (which corresponds to the parameter *μ* in (2.1)) is slightly under-estimated, which results in the estimates of the SNP-level priors and the FDR control being overly conservative. This result is likely because of the combination of our approximation strategy and the relatively small sample size. Utilizing the highly informative quantitative priors substantially improves the power of eQTL discovery (Table 2). For the modest decay rate, incorporating DTSS in eQTL discovery results in a 15% (or 7 percentage points) power gain, whereas in the fast decay case, we find that there is a 25% (or 10 percentage point) increase in power, which results in correctly discovering ~ 1000 more eGenes on average.

**Figure 1:**
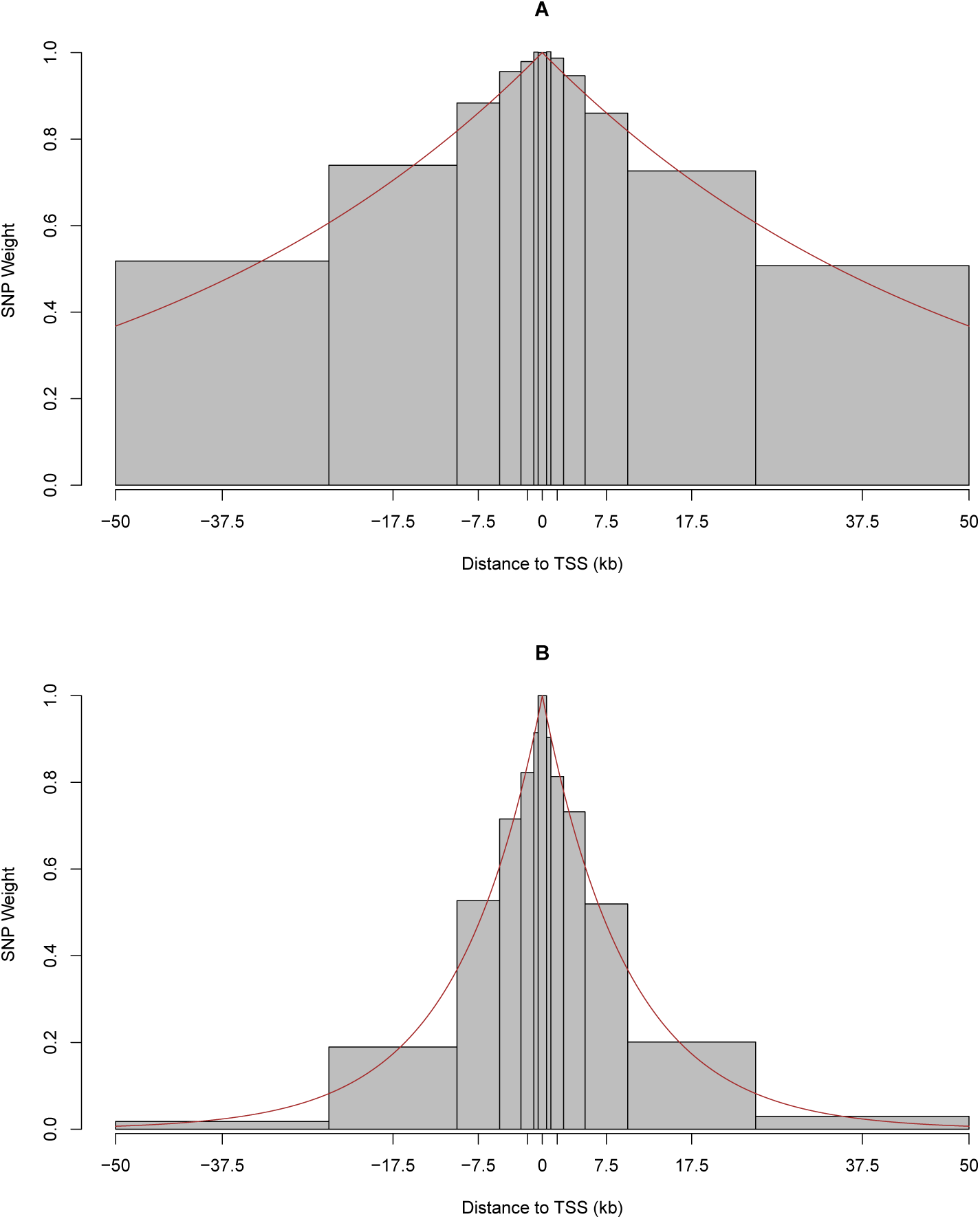
TORUS estimates of eQTL signal decay rates with respect to DTSS in simulations. Panels A and B plot the estimates by the EM algorithm for the modest and fast decay rates, respectively. Each bar in the plot represents a distance bin. To determine the height of the bar, we compute the prior association probability of a SNP located in the corresponding distance bin by plugging in the MLEs (averaged over 20 repetitions) using equation (4.2). We then normalize the resulting probabilities with respect to the center bin such that the center bar always has a weight of 1. For visualization purposes, we choose to highlight the 100 kb region centered around TSS. The red lines in both panels denote the true decay rate according to the generating functions. It is clear that the enrichment estimates from the EM algorithm capture the overall patterns of the decay effect quite accurately.

**Table 2:**
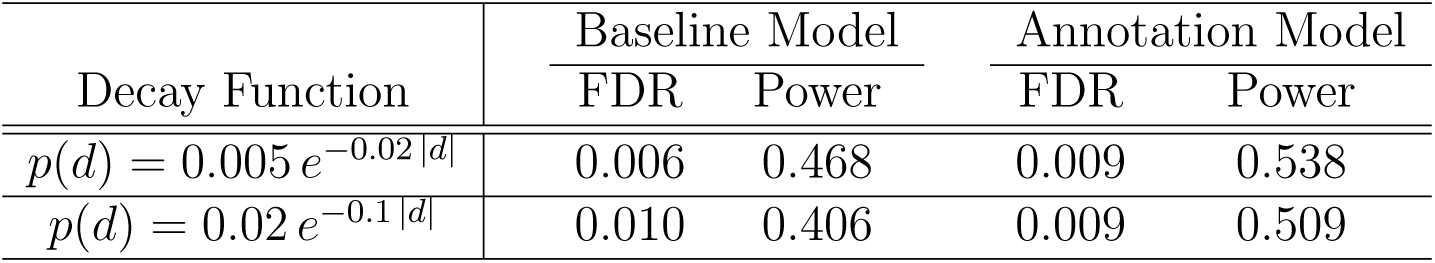
Comparison of QTL discovery with and without incorporating genomic annotations using simulated eQTL data. We simulate the eQTL data sets such that the majority of QTN signals decay according to the function *p*(*d*). The annotation model uses the SNP distance to TSS as annotations, whereas the baseline model does not. For both the modest and rapid decay functions, we observe substantial power gain by incorporating relevant annotations into the QTL discovery.

### 2.3 Analysis of GTEx eQTL Data

We analyze the eQTL data sets from the GTEx project (release version 6), which consist of genotype and expression phenotype data from 44 human tissues. The sample sizes in this data release vary from 70 (uterus) to 361 (muscle skeletal). The genotype and expression data have been subjected to the standard quality control protocols performed by the GTEx consortium. We download the summary-level statistics, 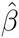, 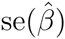, for each gene-SNP pair computed by the software package MatrixEQTL (Shabalin 2012) directly from the GTEx portal. The GTEx portal also provides a list of eGenes for each tissue obtained by the gold-standard minimum *p*-value approach using permutation and Storey’s *q*-value procedure.

We first run the proposed Bayesian method at the baseline without using any annotations to identify eGenes at the FDR 0.05 level, and the result is shown in Figure 2A. Compared with the permutation result, it displays a pattern that is very similar to what we observed in the first simulation study: at the baseline level, the Bayesian method appears to be optimal when the detectable QTL signals are overly low, whereas when the detectable signals are high, it performs slightly worse than the gold-standard approach.

**Figure 2:**
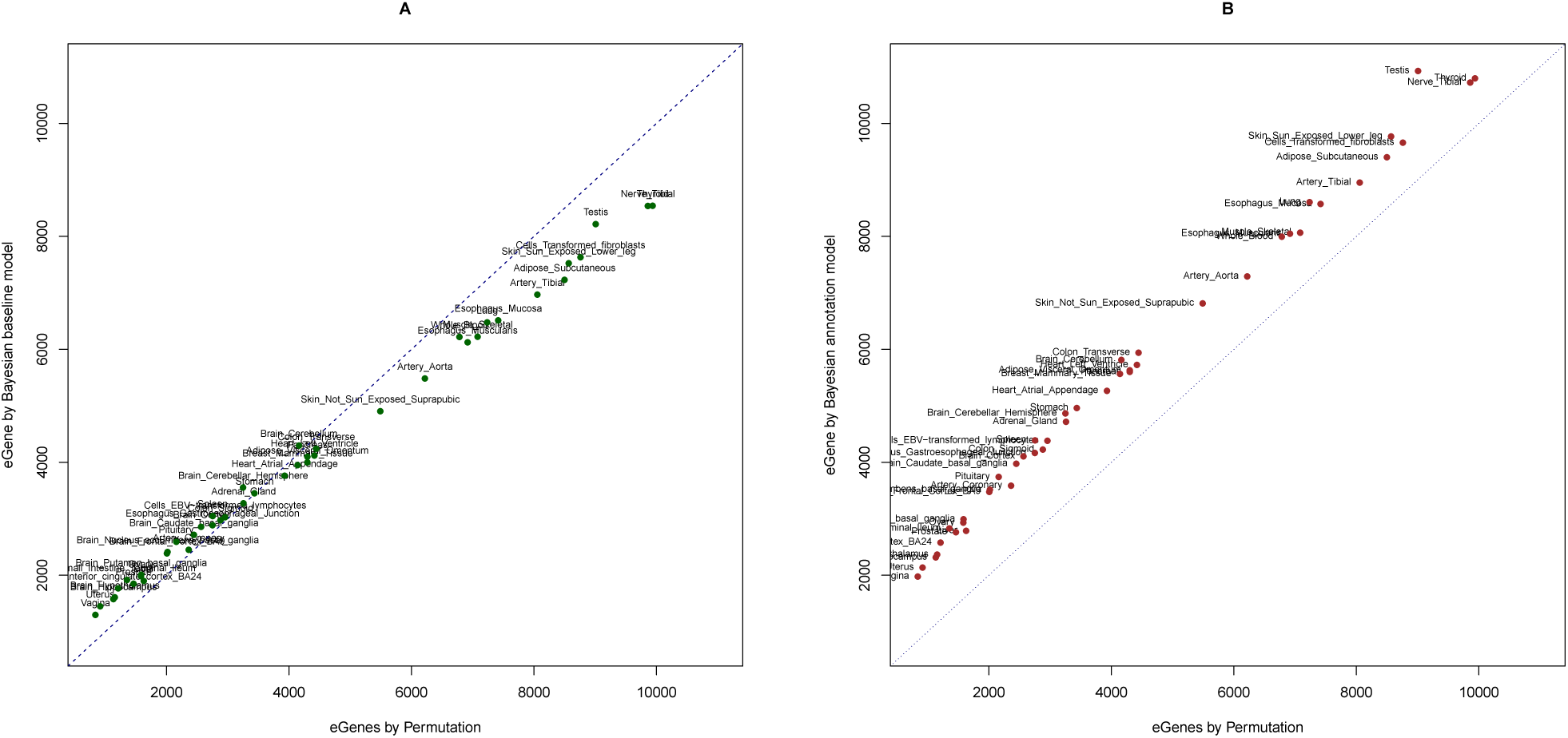
eQTL discovery from GTEx data by TORUS. We plot the number of eGenes discovered by TORUS versus the minimum *p*-value approach in each tissue. Each point represents a single tissue. Panels A and B present the TORUS results in the baseline and by incorporating DTSS annotations, respectively. The pattern observed in panel A is very similar to what we observed in the first simulation study. With the incorporation of DTSS annotations, TORUS discovers more eGenes in all tissues.

We then include the SNP DTSS annotations into the hierarchical model and re-analyze the data using the proposed QTL discovery procedure. We find that the eGene discovery is uniformly improved: in each single tissue, incorporating DTSS yields more eGenes than using either the baseline model or the gold-standard permutation approach. On average, we discover 1,475 more eGenes per tissue compared with the gold-standard approach when incorporating DTSS information in the hierarchical model. Most importantly, we find great concordance between the eGenes discovered: on average, 93% of the eGenes discovered by the gold-standard permutation procedure are also identified by TORUS.

Computationally, both analyses by TORUS complete within 1 hour of running time for a single tissue. On a distributed computing cluster, the full analysis for all 44 tissues takes less than 12 hours.

Our eGene discovery method also yields SNP-level priors incorporating annotations that are critical for the downstream fine-mapping analysis. We perform the multi-SNP eQTL fine mapping for the discovered eGenes in the lung tissue using the MCMC algorithm proposed in Wen *et al*. (2015). We find that in 19 of 8,605 identified eGenes (at the FDR 0.05 level), there is at least one convincing eQTL signal (posterior inclusion probability of > 0.90) that is located 500 kb or farther away from the corresponding TSS. Although relatively rare, the association strength of those signals are quite strong. We take the example of gene *CCZ1* (Ensembl ID: ENSG00000122674) and show the fine-mapping results of its *cis*-eQTLs in Fig. 3. The *cis* region appears to harbor multiple independent eQTL signals, and the posterior inclusion probabilities for three independent signals exceed 99%. The farthest eQTL signal is approximately 900 kb away from the TSS, and the fine-mapping analysis narrows the set of causal variants for this signal to approximately 10 highly correlated SNPs. Through this example, we have illustrated that by defining a relatively large *cis* regulatory candidate region, we are able to find strong eQTL signals that locate farther away from the gene (according to the 1-dimensional map). More importantly, our QTL discovery analysis naturally connects to the downstream fine-mapping analysis by providing the critical SNP-level prior information.

**Figure 3:**
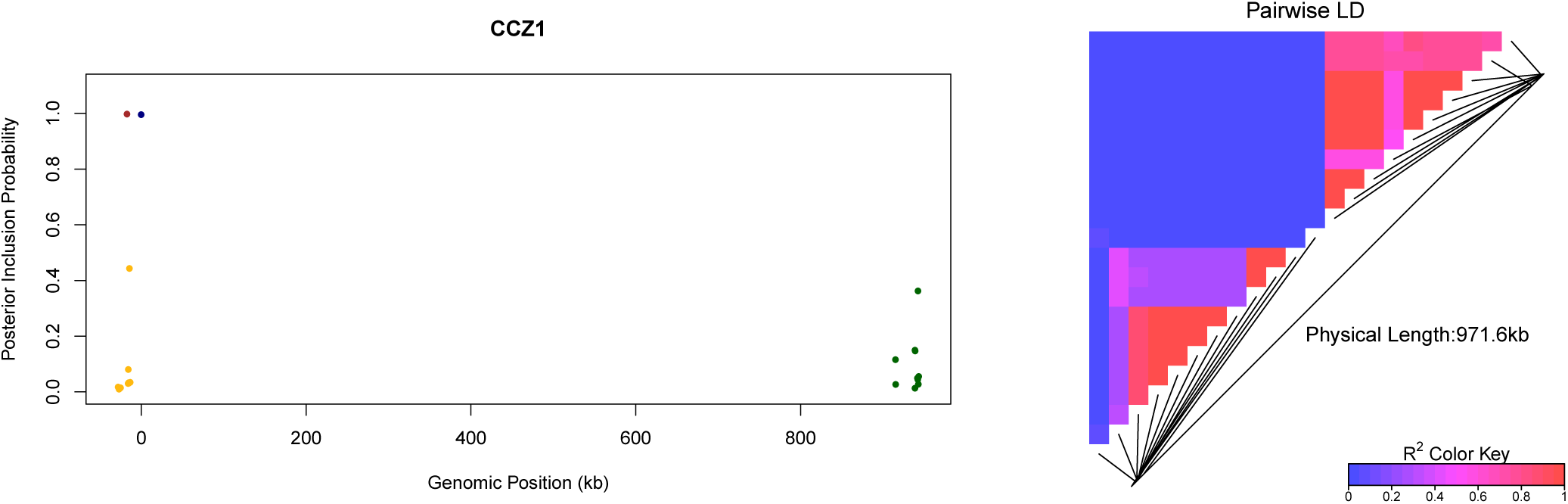
Fine mapping of gene *CCZ1* based on the QTL discovery results in lung tissue. Using the SNP-level prior generated from the QTL discovery analysis, we perform multi-SMP fine mapping using the software package FM-QTL. Overall, we identify 4 clusters of independent association signals. The left panel plots all the SNPs with posterior inclusion probabilities (PIPs) ≥ 0.01. The locations of the SNPs are labeled with respect to the TSS. The SNPs in the same color are in LD and represent the same eQTL signal. The cumulative PIP for the purple, blue and green clusters are all ≥ 0.99, and the cumulative PIP for the gold cluster is 0.72. The heatmap on the right panel represents the LD structure among the plotted SNPs. This analysis demonstrates that some distant eQTL signals (in this case, the green cluster) can be confidently detected, and QTL discovery should include a large candidate region

## 3 Discussion

In this article, we have introduced a powerful statistical approach for discovering molecular QTLs through the use of high-throughput sequencing data and dense genotype data. Through a combination of theoretical derivations, simulation studies and real applications, we have demonstrated that i) our proposed novel approach rigorously controls pre-defined false discovery rates in QTL discovery; ii) by naturally integrating highly informative genomic annotation, the proposed approach consistently displays superior power compared with the current gold-standard approaches; and iii) our implementation of the proposed statistical methods exhibits superb computational efficiency and is one-hundred times faster than the gold-standard method by avoiding extensive permutations.

Our use of hierarchical modeling enables the integration of genomic annotations into QTL mapping in an elegant probabilistic framework. More importantly, it provides an EM algorithm to perform *rigorous* enrichment analysis. Note that the embedded enrichment analysis procedure extends beyond enrichment testing – it provides accurate quantification of the enrichment level of each annotation (Supplementary Material).

The SNP distance to TSS is probably the most convenient genomic annotation. Nevertheless, we have demonstrated that the proper use of DTSS helps to resolve a long-standing dilemma in *cis*-eQTL mapping: the choice of the *cis*-region length. It is well known that most *cis*-eQTL signals are clustered around TSS and become sporadic away from it. This appears to suggest that one should focus on a relatively shorter *cis* region (e.g., ~ 100 kb) to lessen the multiple testing burden and discover more eGenes. However, such an approach will inevitably miss some distant yet strong signals, and the accumulative loss of signals across all genes can be severe. In our proposed approach, we select a rather large *cis* region and use the enrichment analysis to assess a prior weight of each SNP by their DTSS. Consequently, neighboring SNPs of TSS are up-weighted, and distant SNPs are relatively down-weighted. This naturally solves the dilemma: the focus is on close-by SNPs, but strong distant signals can still overcome the prior weighting penalty and be uncovered.

Finally, in our view, QTL discovery is not the endpoint of the QTL analysis. Rather, it primarily serves as a screening procedure to prioritize a subset of candidate loci that are highly likely to harbor causal trait-associated variants – a strategy that is well demonstrated and widely applied in genome-wide association studies (GWAS). As illustrated in our analysis of GTEx data, our QTL discovery approach naturally connects to the downstream (Bayesian) multi-SNP fine-mapping analysis by supplying the SNP-level priors computed by plugging in the point estimate from the enrichment analysis into the prior model (4.2).

## 4 Methods

### 4.1 Hierarchical Model for QTL Discovery

We consider a general problem of QTL mapping at the genome-wide scale. In particular, we assume that there are *L* genomic loci (in many cases, the loci are naturally formed by genes), each of which contains *p_l_* SNPs for *l* = 1, …, *L*. Given a sample of *n* unrelated individuals, for each locus *l*, we model the potentially locus-specific quantitative trait measurement ***y_l_*** within the sample using the following general form of the linear regression model

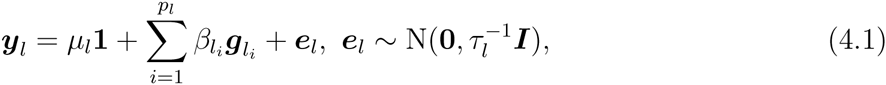

where *n*-vectors 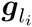, ***e*** represent the sample genotypes at genetic variant *l_i_* and the residual errors, respectively. (At present, we assume that ***y*** is a univariate quantitative trait, and we relax this assumption and extend the framework to multivariate quantitative traits in section 4.4.) Furthermore, we denote 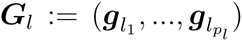. Note that the model allows multiple SNP associations within a given locus. The regression coefficients *μ_l_* and 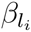 represent the intercept and the genetic effect of each genetic variant, respectively, and 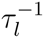 denotes the residual error variance. Following Wen (2014), we further denote the latent binary association status of each genetic variant *l_i_* by 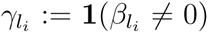 (i.e., 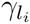 is dichotomized from the corresponding 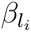), and 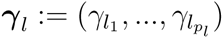.

Our prior specifications for the parameters *μ*, 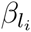 and τ are mostly standard and follow directly from Marin and Robert (2007), and we leave the details to the Supplementary Materials. Most importantly, we use the prior specification for *γ_l_* to incorporate variant-level genomic annotation information. Specifically, we assume that the 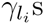 are *a priori* independent and that

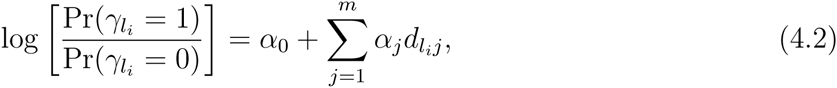

where we use 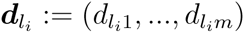 to denote the variant-specific genomic features, and the hyperparameter ***α***:= (*α*_0_,…, *α_m_*), which characterizes the enrichment level of each genomic feature in trait-associated genetic variants, is referred to as the enrichment parameter. Note that in the special case where no genomic annotation is used in the analysis, the prior model (4.2) contains a single parameter *α*_0_, which quantifies the prevalence of trait-associated genetic variants among all candidate SNPs. We refer to this model as the baseline model.

### 4.2 Multiple Hypothesis Testing and Bayesian FDR Control

The problem of QTL discovery can be framed as a hypothesis testing problem. Specifically, we identify locus *l* as a QTL if the null hypothesis asserting that it contains no trait-associated genetic variant, i.e.,

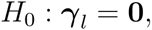

is rejected. The problem of multiple hypothesis testing arises because we perform simultaneous testing for tens of thousands of loci across the genome when mapping molecular QTLs.

To take full advantage of our hierarchical model, we adopt a Bayesian FDR control strategy (Newton *et al*. 2004; Müller *et al*. 2004). Briefly, the Bayesian FDR control requires computing the posterior probability *u_l_*:= Pr(*γ_l_* = 0 | ***y****_l_*, ***G****_l_*) for each locus *l*, and it rejects the null hypothesis if *u_l_* is small (note that the rejection rule is analogous to the commonly applied *p*-value-based approach). Based on a pre-defined FDR control level *α*, a straightforward algorithm (Newton *et al*. 2004) can be applied to determine the induced rejection threshold *t_α_* such that

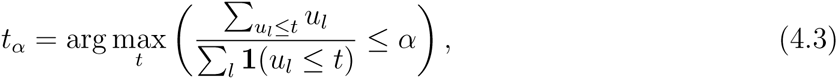

where the expression 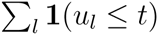 in the denominator represents the total number of rejections at threshold *t*, and the expression 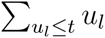 in the numerator represents the expected false rejections at threshold *t*. The Bayesian FDR control procedure is naturally connected to its frequentist counterpart (Supplementary Material), with the primary difference being that the Bayesian FDR is conditional on the observed data in hand whereas the frequentist procedure computes FDR over hypothetically repeated experiments. Furthermore, Müller *et al*. (2004) proved that the Bayesian procedure is theoretically optimal in the sense that it minimizes the corresponding false non-discovery rate (FNR, a measure of power).

### 4.3 Approximate Computation of Posterior Probability

Although the posterior probability *u_l_* is conceptually straightforward to obtain from the proposed hierarchical model, its computation is practically intractable. To ease the computation, we apply two levels of approximation.

A critical intermediate step in evaluating *u_l_* is to compute the probability Pr(***γ* = 0 | *y_l_, G_l_, α*)** for a given value of the enrichment parameter ***α***. Specifically, we evaluate this quantity by

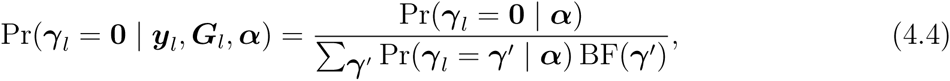

where BF(*γ*) denotes the Bayes factor

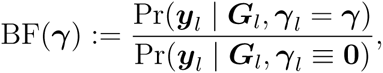

and by definition, BF(**0**) = 1. Although the calculation of BF(*γ*) for any given *γ* can be achieved analytically for a wide range of linear model systems (Wen 2014), it is practically infeasible to enumerate all possible *γ* values when the number of SNPs within a locus is large (for *p* SNPs in a locus, there are a total of 2*^p^* association models to explore). Here, we propose using the approximation

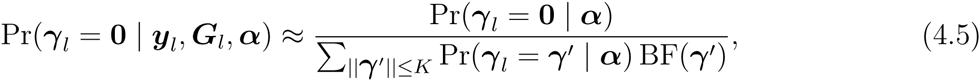

where 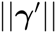 denotes the number of non-zero indicators in vector *γ*′, and the subset 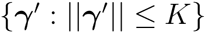 consists of only the association models with no more than *K* associated SNPs. Note that the approximation (4.5) is practically accurate if

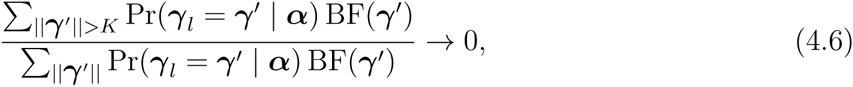

i.e., the posterior probability on more complicated association models becomes negligible. Nevertheless, even if this ideal condition is not satisfied, the approximation still provides a *conservative* estimate of the true posterior probability because

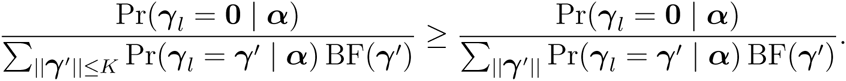

This property is critically important for ensuring that applying approximation (B.2) does not increase type I errors.

To achieve the best computational efficiency, we simply set *K* = 1 in practice, which is optimal when there is at most one causal SNP per locus. In this special case, the approximation has aanalytic form, i.e.,

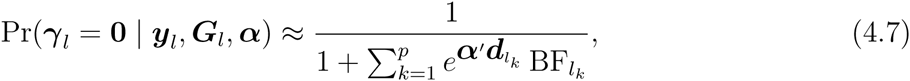

where 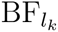 denotes the Bayes factor for the *k*-th SNP from the single SNP testing and can be computed analytically from the corresponding single SNP testing statistics (Servin and Stephens 2007; Wakefield 2009). One of the most attractive advantages is that the full analysis now only requires summary-level statistics rather than full individual-level genotype-phenotype data. Note that the assumption of ”one QTN per locus” is widely used in genetic association/QTL mapping analyses. The gold-standard frequentist test statistic, the minimum *p*-value in a locus by single SNP association testing, is the most powerful if only one genuine association exists within the locus (De la Cruz *et al*. 2010). Many Bayesian approaches make this assumption explicitly (Veyrieras *et al*. 2008; Gaffney *et al*. 2012; Flutre *et al*. 2013; Pickrell 2014) in prior specification. Although similarly intuitive, restricting the number of causal variants *a priori* requires a completely different formulation that is not as straightforward to interpret (as our logistic priors) and often causes convergence issues when estimating the enrichment parameters. Moreover, we argue that for most of the currently available molecular QTL data sets, we only discover one convincing association signal for the vast majority of the quantitative trait loci (Wen *et al*. 2015). Therefore, we expect that this approximation should be accurate on average with no severe loss of power.

Additionally, we take an empirical Bayes approach to estimate ul by

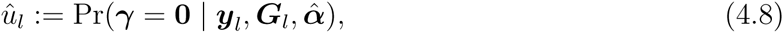

where 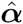 denotes the maximum likelihood estimate of the enrichment parameter. This approach essentially replaces the integration (of ***α***) by a more computationally efficient optimization procedure. To find the MLE of ***α***, we follow (Wen *et al*. 2015) and derive an efficient EM algorithm that incorporates the analytic posterior approximation of Pr(γ*_l_* | ***y_l_***, ***G_l_***, ***α***) (details are described in the Supplementary Material). Briefly, in the E-step, we compute the posterior inclusion probability (PIP) of each SNP in each locus given the current estimate of ***α***, and in the M-step, we simply fit a logistic regression model using the PIPs as the response variable and update the estimate of ***α*** by the corresponding fitted regression coefficients. Note that the PIP computation in the M-step also applies the posterior approximation that essentially assumes a single QTN per locus. Interestingly, through extensive simulations (Supplementary Fig. 1), we find that the intercept term α_0_ in (4.2) tends to be slightly under-estimated. This result is partially due to the conservative nature of our posterior probability approximation, but it is mostly related to the imperfect power to detect associations, i.e., QTNs with small effects are difficult to identify with a limited sample size even when a multi-SNP model is considered. Interestingly, other enrichment coefficients associated with specific annotations are generally estimated unbiased and accurately (Supplementary Fig. 1). Overall, the prior estimates from the EM algorithm are slightly conservative, which is arguably desirable in QTL mapping problems.

### 4.4 Extension to Multivariate Quantitative Traits

Thus far, our description of the proposed method has focused on univariate quantitative traits, e.g., gene expressions and DNA methylation measurements. Our framework can be trivially extended to applications in which the quantitative trait is measured by multivariate variables, e.g., in the case of using ATAC-seq data to quantify chromatin accessibility. To accommodate multivariate quantitative trait data, we simply replace the model (4.1) by a multivariate linear regression model, which naturally accounts for the correlations between multiple components of the trait. In the example of ATAC-seq data, the response variable for each individual at each locus can simply be described by a row vector with each entry representing the read counts from a pre-defined window on the genome. To perform the Bayesian FDR control, it only requires to adjust the single SNP association Bayes factor according to the modified multivariate linear regression model. Fortunately, such results are available in the statistical literature (Wen and Stephens 2014).

### 4.5 Extension to QTL Data Composing Multiple Heterogeneous Groups

Molecular QTL data collected from multiple heterogeneous sources have become increasingly available (Maranville *et al*. 2011; Barreiro *et al*. 2012; Wen *et al*. 2015; Ardlie *et al*. 2015). Joint analysis of QTL data across multiple heterogeneous groups not only improves the power of identifying consistent QTL signals across groups (Flutre *et al*. 2013; Wen *et al*. 2015) but also helps to correctly map group-specific QTL signals (Maranville *et al*. 2011; Barreiro *et al*. 2012; Flutre *et al*. 2013). Utilizing the previous statistical results from computing Bayes factors from heterogeneous subgroups (Wen and Stephens 2014), the proposed approach can be trivially applied in those scenarios for QTL discovery while integrating genomic annotations.

## 5 Data Access

The GTEx summary-level statistics can be downloaded from the GTEx portal (http://www.gtexportal.org/home/). The GEUVADIS data are publicly available at http://www.geuvadis.org/web/geuvadis/RNAseq-project. The simulation scripts and the software package TORUS (including source code) can be downloaded from https://github.com/xqwen/torus/

## Appendix A Bayesian FDR Control and Its Connection to Frequentist Approaches

In the context of QTL discovery, the multiple testing problem can be framed as a binary decision problem with respect to *γ_l_* for *l* = 1,…, *L*. We define a binary indicator, *Z_l_*, for each locus *l* and set *Z_l_* = 1 if *γ_l_* = 0 and 0 otherwise. Further, we denote the collection of the observed phenotype-genotype data by ***Y***. Let the function *δ_l_ (****Y***) denote a decision (0 or 1) on *Z_l_* based on the observed data, and define the total discoveries by 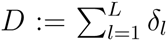. Following the formulation of Müller *et al*. (2006), the False Discovery Proportion (FDP), which is also a random variable, can be defined as the proportion of false discoveries among total discoveries, i.e.,

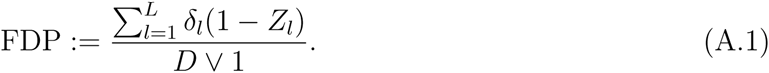

Recall,

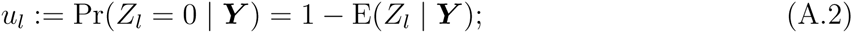

thus, the Bayesian False Discovery Rate is naturally defined as

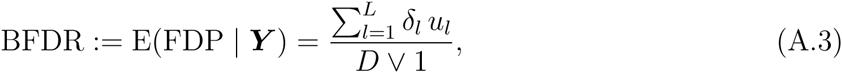

where the conditional expectation is taken with respect to ***Z***:= (*Z*_1_,…, *Z_L_*). Moreover, the frequentist control of the False Discovery Rate focuses on the quantity

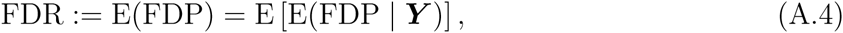

where the additional expectation is taken with respect to ***Y*** over (hypothetically) repeated experiments. It is important to note that controlling the Bayesian FDR is a *sufficient but not necessary* condition to control the frequentist FDR; thus the Bayesian FDR control is more stringent in theory.

As demonstrated by Newton *et al*. (2004) and Müller *et al*. (2006), the Bayesian FDR control is based on the following natural decision rule

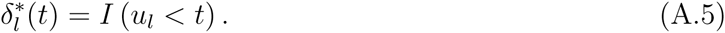

For a pre-defined FDR level *α*, the threshold *t_α_* in (A.5) is determined by

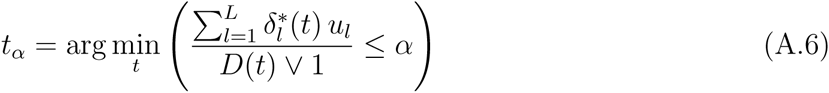

In practice, we use the following simple algorithm proposed by (Newton *et al*. 2004) to determine *t_α_*:

1. *sort u_l_’s in ascending order: i.e*., 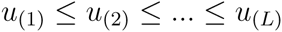
2. *start from m* =1 *and compute the partial mean using the sorted sequence of* {*u*_(_*_l_*_)_} *for* 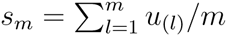
3. *stop if s_m_ > α*
4. *t_α_* — *u*_(_*_m–1_*_)_ *if m > 1, and 0 otherwise; and reject the hypotheses corresponding to u*_(1)_, *…,u*_(_*_m–1_*_)_.

## Appendix B EM Algorithm for Enrichment Analysis

### B.1 Algorithm Details

In this section, we outline the EM algorithm to estimate the enrichment parameter ***α***. The algorithm is a special case of what is described in Wen *et al*. (2015). We denote 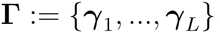. By treating **Γ** as missing data, we obtain the complete data likelihood by the following

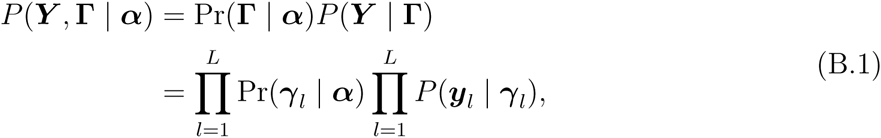

where the factorization is based on the conditional independence relationships induced by the hierarchical model. We further re-write the prior probability Pr(*γ_l_* | ***α***) using the logistic model,

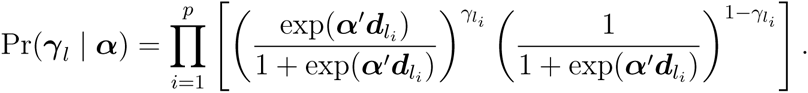

Therefore, the complete data log-likelihood is given by

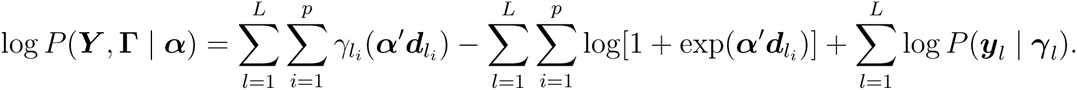

The EM algorithm is initiated at an arbitrary starting point ***α*** = ***α***^(1)^. In the E-step of the *t*-th iteration, we compute

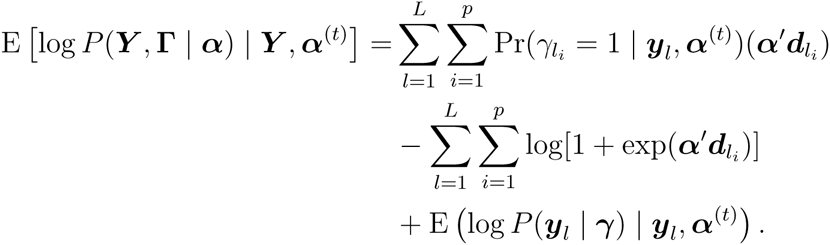

That is, we evaluate the posterior inclusion probability 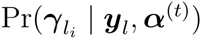 for each candidate SNP and for all loci. In the M-step, we find

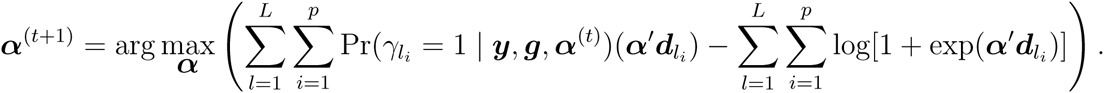

It is not difficult to recognize that the the functional form of the objective function coincides with the log likelihood function of a logistic regression model with the binary response variable, 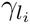, replaced by its corresponding posterior expectation, 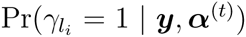. Therefore, we can directly fit a logistic regression model to find ***α***^(^*^t^*^+1)^.

The computational difficulty of the EM algorithm lies in evaluating the PIPs in the E-step. The situation is the same as computing *u_l_* in Bayesian FDR control where the exact computation is intractable. To ease computation, we apply the same deterministic approximation technique. The key assumption is again that posterior probabilities of single QTN association models dominate the posterior probability space of {***γ***} for locus *l*, i.e.,

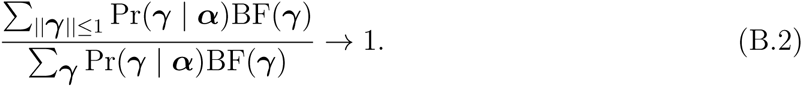

Consequently, it follows that

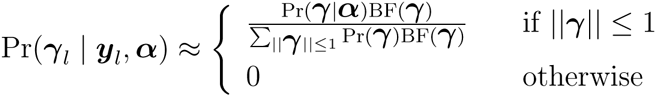

The model space of 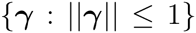 contains only the null model, *γ_l_* = 0, and all single-SNP association models. We denote

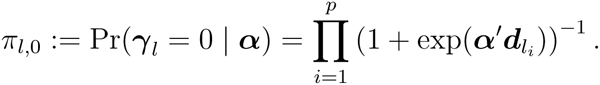

We use 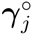 to denote the single-SNP association model where the *j*-th SNP is the assumed QTN. Clearly,

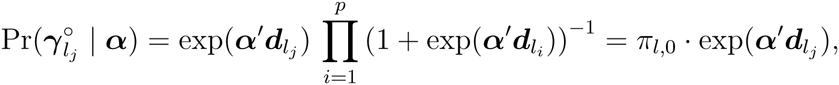

and

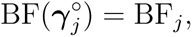

and recall that BF*_j_* denotes the Bayes factor based on the single-SNP analysis of SNP *j*. Finally, we note that given the restrained model space, the PIP of SNP *j*, 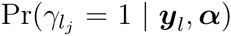, coincides with the posterior model probability, 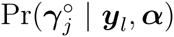. Given all of the above, it follows from the simple algebra that

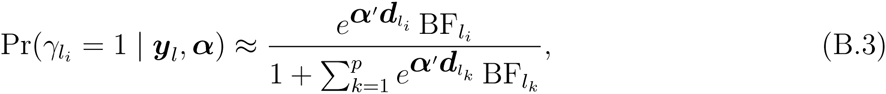

which can be analytically evaluated given ***α***.

### B.2 Accuracy Evaluation by Simulation

We perform simulation studies to evaluate the performance of the EM algorithm in enrichment analysis. Our simulation setting mimics the application of genome-wide *cis*-eQTL mapping, however at a reduced scale. Specifically, we select a subset of 5,000 random genes from the GEUVADIS data. For each gene, 50 *cis*-SNPs are used in the simulation and we annotate 30% of the SNPs with a binary feature. For each SNP, the association status is determined by a Bernoulli trail with the success (i.e. associated) probability given by

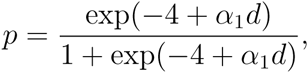

where *d* is the SNP specific binary annotation value, and α_1_ is the true enrichment parameter. Given the true QTNs of each gene, we then apply the scheme described in section C to simulate the effect sizes of the QTNs and the expression levels for the 343 European individuals. We set *α*_1_ = −0.25, 0.00, 0.25, 0.50, 0.75,1.00, and for each *α*_1_ value, we simulate 100 data sets. We use the proposed EM algorithm to analyze the simulated data sets. For comparison, we also estimate *α*_1_ using a logistic regression with the true association status as the outcome variable and the annotations as the predictor. This analysis represents a theoretical best case scenario, and its results should be regarded as the optimal bound for the analyses that infer the latent association status from the genotype-phenotype data. The results (Supplementary Fig. 4) indicate that the EM algorithm based on our posterior approximation scheme consistently yields unbiased estimates for *α*_1_. The decrease of estimation efficiency from the theoretical optimal estimator, represented by the difference of the standard errors, is not large. However, we do find the baseline parameter *α*_0_ is consistently under-estimated, largely due to imperfect power to detect associations.

**Figure 4:**
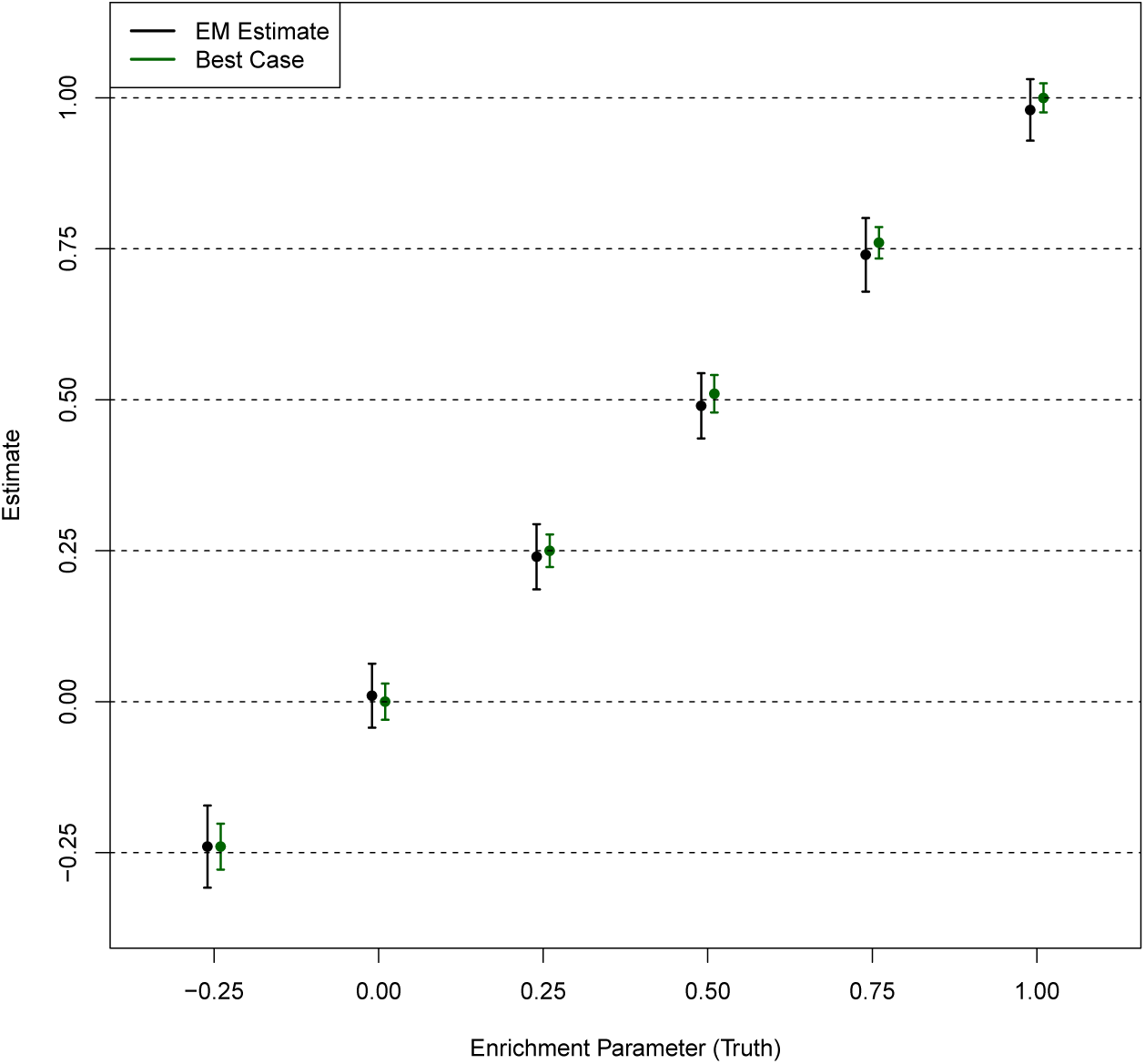
Point estimates of the enrichment parameter in simulations. The point estimate of *α*_1_ ± standard error (obtained from 100 simulated data sets) for each method is plotted for each simulation setting. The “best case” method uses the true association status and represents the optimal performance for any enrichment analysis method. The estimate based on the EM algorithm using the posterior approximation yields unbiased estimate but with larger variation than the optimal method, which is fully expected.

## Appendix C Simulation Details

In this section, we provide the details of the scheme for simulating genetic effects of casual QTNs and the individual-level quantitative traits.

As described in the main text, we perform Bernoulli trials for each of the candidate SNPs and determine its association status with the target (expression) quantitative trait. For each causal QTN, we then draw its genetic effect from a Normal distribution, i.e.,

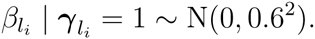

The individual-level expression levels for locus *l* are then simulated according to the linear model

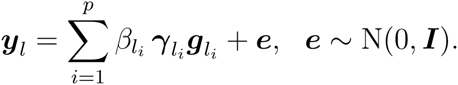

In our scheme, the genetic association by a causal QTN explains 0.7% (for QTN with minor allele frequency 1%) to 15% (for QTN with minor allele frequency 50%) of the heritability, which is quite realistic.

In the analysis, we compute the single SNP Bayes factor using the analytic formula by Wakefield (2009). More specifically, we assume the prior genetic effect of an QTN is drawn from a mixture of Normal distribution, i.e.,

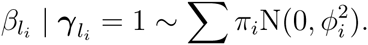

And we use a grid of *ϕ* values, (*ϕ*: 0.1, 0.2, 0.4, 0.8}, and set *π*_1_ = ⋯ = *π*_4_ = 0.25. We also apply this setting for analyzing the eQTL data from the GTEx project.

## Appendix D Binning of *cis*-SNPs by DTSS

Here we describe the binning scheme to annotate the SNPs in the *cis* region of a target gene. We place each SNP into 21 unequally spaced bins according to its DTSS (Table 3). The the bins are smaller and denser as close to TSS, which helps capture the rapid decay of QTL signals. This binning scheme leads to estimation of 21 enrichment parameters in the enrichment analysis.

**Table 3:** Binning scheme for *cis*-SNPs

| Bin | Range (kb) | Size (kb) |
| --- | --- | --- |
| -10 | $< -500$ | 500 |
| -9 | $[-500, -250)$ | 250 |
| -8 | $[-250, -100)$ | 150 |
| -7 | $[-100, -50)$ | 50 |
| -6 | $[-50, -25)$ | 25 |
| -5 | $[-25, -10)$ | 15 |
| -4 | $[-10, -5)$ | 5 |
| -3 | $[-5, -2.5)$ | 2.5 |
| -2 | $[-2.5, -1)$ | 1.5 |
| -1 | $[-1, -0.5)$ | 0.5 |
| 0 | $[-0.5, 0.5)$ | 1 |
| 1 | $[0.5, 1)$ | 0.5 |
| 2 | $[1, 2.5)$ | 1.5 |
| 3 | $[2.5, 5)$ | 2.5 |
| 4 | $[5, 10)$ | 5 |
| 5 | $[10, 25)$ | 15 |
| 6 | $[25, 50)$ | 25 |
| 7 | $[50, 100)$ | 50 |
| 8 | $[100, 250)$ | 150 |
| 9 | $[250, 500)$ | 250 |
| 10 | $> 500$ | 500 |

